# *DomainRank*: Improving Biological Data Sets With Domain Knowledge and Google’s PageRank

**DOI:** 10.1101/2022.09.26.509059

**Authors:** Michael Schneider, Juri Rappsilber, Oliver Brock

## Abstract

**Motivation:** The quality of biological data crucially affects progress in science. This quality can be improved with better measurement devices, more sophisticated experimental designs, or repetitious measurements. Each of these options is associated with substantial costs. We present a simple computational tool as an alternative. This algorithmic tool, called *DomainRank*, leverages simple domain knowledge and overlapping information content in biological network data to improve measurement quality at a negligible cost. Following the simple computational template of *DomainRank*, researchers can boost the confidence of their own data with little effort.

**Results:** We demonstrate the performance of *DomainRank* in three test cases: *DomainRank* finds 14.9% more interactions in quantitative proteomics experiments, improves the precision of predicted residue-residue contacts from co-evolutionary data by up to 11.6% (averaged over 882 proteins), and identifies 89.2% more cross-links in photo-crosslinking/mass spectrometry (photo-CLMS) experiments. Although our proposed template is specialized on biological network data, we view this approach as an universal computational tool for data improvement that could be routinely applied in many disciplines.

**Availability:** An implementation of *DomainRank* is freely available: https://github.com/Rappsilber-Laboratory/pagerank-refine

## Introduction

The quality of data greatly affects a scientist’s ability to obtain scientific insight. Any improvement to the quality of scientific data promises to also facilitate such insight. We present a widely applicable, simple, purely computational method for improving the quality of measurement data, without the need for taking additional measurements or acquiring improved measurement devices. The only condition for this to work is that the information contained in subsets of measurements has to exhibit a known relationship.

The intuition is as follows. Let us assume that we have obtained many measurements about a single biological system. These measurements are inherently uncertain. Let us further assume that subsets of these measurements are not independent, i.e. they contain corroborating or complementary information about a particular aspect of the measured system. If we are able to identify and combine this corroborating information, we can use it to improve the quality of the respective measurement data. To achieve this, the main computational challenges are *1)* to identify corroborating information across different measurements and *2)* to use this information to improve the data obtained from the measurements. For the first challenge, we demonstrate that basic domain knowledge suffices. For the second challenge, we show that Google’s PageRank algorithm [34] can leverage this information to achieve substantial quality gains for the data. Together, these two components represent an algorithmic template that can easily be adapted to different types of data and problem domains. We refer to this algorithmic template as *DomainRank*, a portmanteau of domain knowledge and PageRank.

We evaluate *DomainRank* in three different biological domains: protein-protein interactions, residue-residue contacts predicted from evolutionary data, and photocrosslinking/mass spectrometry (photo-CLMS). The focus of our experiments—and this is important to note—is *not* in making a contribution to one of these three domains but instead to demosntrate the general applicability of the proposed algorithmic template. The effectives of *DomainRank* must be measured in relative improvements of data quality. Taking this as a quality criterion, we show that *DomainRank* improves the data in all of these application domains: *DomainRank* identifies 14.9% more interactions from proteomics experiments than the original approach [22], improves the precision of predicted residue-residue contact data [29] by 6.7-11.6% (averaged over 882 proteins), and identifies 89.2% more cross-links compared to the original scoring method [2]. The generality of *DomainRank* and the ease of adapting it to a particular domain enables researchers to boost the quality of their data in their own domain.

## Related Work

PageRank is often used to analyze biological data when it is already represented as a network, i.e. the relationship between different measurements is known. In this section, we survey applications of PageRank to different types of such networks, grouped by the type of data they have been applied to. The key difference between the proposed method *DomainRank* and the related work is the following. Existing methods improve *known* networks. The proposed algorithmic template *DomainRank* leverages domain knowledge to create *novel* networks and then applies PageRank. This means that *DomainRank* opens up the use of PageRank to biological data where a network structure is not known a priori but can be created from available domain knowledge.

### OMICS data

Data from OMICS experiments is noisy, making it difficult to draw biological conclusions. Combining OMICS data sets (for example gene expression data) with network data can improve their reliability, which in turn might lead to novel biological insight. Researchers used such network-based methods to improve target identification by leveraging metabolic data [1, 15], protein-protein interaction data [25], or gene ontology, and expression profile correlations [31]. Further studies use network-based analysis to identify functional complexes in protein-protein interaction networks [12, 40], to refine the protein-protein interaction networks themselves [23, 19, 3], and to predict the outcome of cancer therapy [48].

Some studies use PageRank for target identification. One approach is to construct a network out of secondary data (such as gene ontology) and to re-rank the initially ranked targets [31, 15, 48]. This is a fairly general use of PageRank that is applicable as long as information for constructing the network is available. The studies by Li and Singh et al. [25, 40] use PageRank variants to mine information across heterogeneous networks or protein-protein interaction networks from different species. Although these studies demonstrate only one specific application, these methods could be generalized to other domains. Other studies use modifications of PageRank that are more specific to the problem domain. Banky et al. [1] use a modified PageRank to lower the overwhelming influence of “hubs” in metabolic networks to rank low-degree targets that might be physiologically relevant. Hanna et al. [12] use a pruning procedure that is specific to protein-protein interaction networks. Our proposed method is on the general side of this spectrum, because we demonstrate in this study that we can improve interactions from three different problem domains.

These works discussed above mine information in biological networks, but not to improve the interactions of the networks themselves. Some groups set out to improve the interaction networks [23, 19, 3], but none of these methods use PageRank. We also notice that the study by Kamburov et al. [19] uses a “link graph” representation that is similar to our corroborating information graph. However, we integrate additional information sources from domain knowledge for network refinement while Kamburov et al. use only topology to refine the network [19].

### Residue-residue contact data

Residue-residue contacts are residue pairs that are spatially close in the 3D structure of a protein. These contacts are a powerful information source that can guide protein structure prediction [49, 21, 44, 32, 30]. Many contact prediction methods that have been developed in the recent years [20, 28, 17, 18, 38, 47, 41, 6, 46] use a refinement step to improve their prediction accuracy [5, 41, 18, 47, 46]. This refinement step is often performed by a stacked machine learning classifier [5, 41, 18, 46]. These refinement methods are usually trained on the outputs of the lower-level classifiers and therefore highly specific. In contrast, *DomainRank* is able to refine contact maps from different methods and does not require training on specific lower-level methods. Therefore, our proposed template is more general than the related approaches discussed here. Additionally, our method can be applied to other problem domains.

Frenkel-Morgenstern et al. [8] developed a networkbased method. This method encodes predicted contacts into a graph data structure (residues as nodes and contacts as edges) and performs edge clustering to filter contacts. In contrast, we use a different graph representation (contacts as nodes, corroborating information as edges) and additional information to refine contacts.

## Methods

### The *DomainRank* Template

We now give an overview of *DomainRank* and how it can improve interactions in biological network data (Fig. 1). These noisy interactions may come from different sources and contain true and false positives. We use corroborating information between interactions to construct a *corroborating information graph* and use PageRank [34, 24] to rescore the interactions to leverage this corroborating information. PageRank propagates corroborating information through the graph until a steady state is reached. In this steady state, the score of a node (representing an interaction) is now proportional to the amount of corroborating information that is pointing to that node. This procedure re-scores the input interaction, effectively increasing the confidence of interactions that are highly supported by corroborating information.

**Figure 1:**
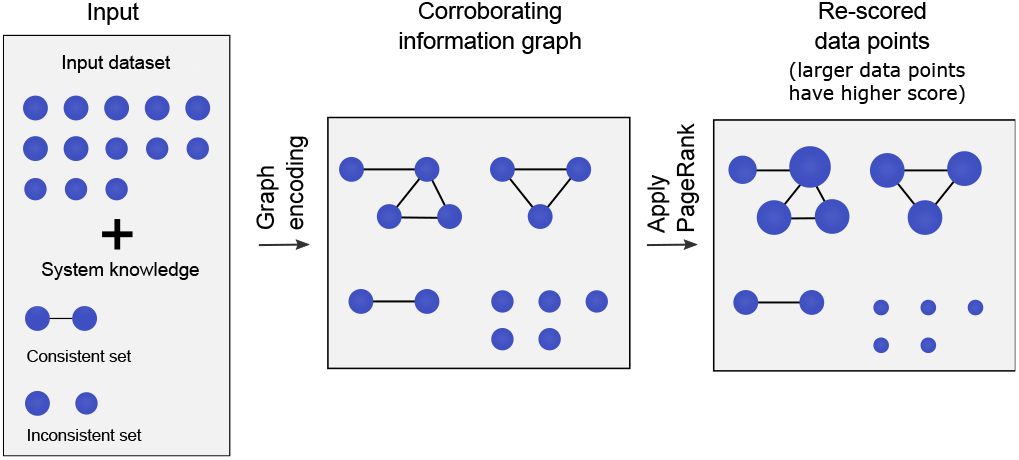
Outline of our proposed computational template. A data set (containing true positive and false positive data points) and prior knowledge of the observed system is the input. We then build the corroborating information graph in which two data points are connected by an edge if their consistency is supported by the system structure. We then use Google’s PageRank algorithm to propagate corroborating information through the corroborating information graph. This assigns higher scores to data points that are well supported by corroborating information (larger data points indicate higher scores).

The general template consists of two steps: 1) Defining the corroborating information between interactions from knowledge about the problem domain and 2) using PageRank for re-scoring. Note that step two is influenced by two parameters (see section Methods). The first parameter α balances the relative importance of the initial confidence in the input interactions and the confidence derived with corroborating information. The second (optional) parameter is the number of interactions used to construct the corroborating information graph. When too many low-confidence input interactions dominate the corroborating information graph, artificial clusters might lead to high scores of wrong interactions, which leads to deteriorated performance.

In the Results and Discussion section, we provide a high-level overview of the corroborating information for our three application domains. Details of how the algorithmic template can be adapted for these three domains is includined in the Supplementary Materials.

We applied DomainRank to three biological network data sets: A protein-protein interaction (PPI) data set from quantitative proteomics experiments, predicted residueresidue contact pairs from co-evolution data, and high-density cross-linking/mass spectrometry data.

### Refinement of Protein-Protein Interaction Data

Protein-protein interactions mediate many biological processes and therefore play a central role in our understanding of the cell. Because of their importance, researchers developed many high-throughput methods to measure these protein-protein interactions, such as proteome fractionation [13], yeast-two hybrid screening [42], affinity purification [9], and crosslinking/mass spectrometry [35]. Regardless of the underlying method, protein-protein interaction data contains a significant number of false positive interactions. Thus, algorithms that reduce these false positives increase the biological interpretability of this data or increase the “depth” of the measurement, which is the number of protein-protein interactions that can be detected.

Here, we used *DomainRank* to improve protein-protein interaction data from quantitative mass spectrometry experiments [22]. We used Gene Ontology (GO) information to construct the corroborating information graph. We assume that two protein-protein interactions in similar subcellular environments corroborate each other because the environment facilitates these interactions (see Methods for details). This is also in line with earlier studies by Morrison et al., which found that networks based on subcellular information are effective to rank gene expression changes from microarray experiments [31].

We tested whether *DomainRank* can increase the number of detected protein-protein interactions from this mass spectrometric analysis. We analyzed the chromatogram data from the original paper following the steps described in the original study and used this data as input for our method [22]. *DomainRank* increases the number of protein-protein interactions by 14.9% (absolute increase from 9886 to 11360 interactions) at the same target precision of 0.53 used in the original study (Fig. 2) [22]. At a precision cutoff of 0.64 or higher, the original data analysis method and *DomainRank* return approximately the same number of interactions, which suggests that *DomainRank* especially boosts low-confidence interactions. Note that the absolute numbers are not directly comparable to the original study, because we used a newer version of the CORUM database for validation (accessed December 2016) [37].

**Figure 2:**
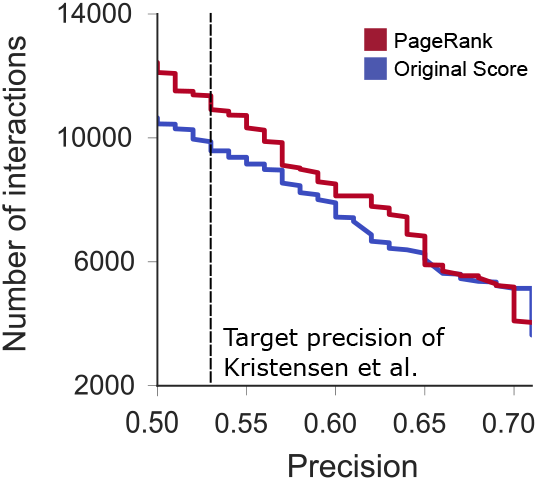
*DomainRank* improves the number of interactions from quantitative mass spectrometry experiments. At the same precision cutoff of 0.53 used in the original study [22], *Domain-Rank* identifies 14.9% more interactions then the original data analysis protocol. The increase is most pronounced at low precision cutoffs, which suggests that *DomainRank* especially boosts low-confidence interactions.

In summary, our template increases the number of protein-protein interactions identified by quantitative mass spectrometry with a fast and simple post-processing step. We believe that *DomainRank* can also improve data from other high-throughput methods. We would like to state again that we do not make any claims about advancing the field of refining protein-protein interaction data; we only demonstrate that DomainRank can be applied beneficially to this type of data.

### Refinement of Predicted Contact Data

Residue-residue contact prediction methods predict residue pairs that are in spatial proximity in the folded protein. Predicted residue contacts can be the decisive factor for successful *ab initio* structure prediction [32]. Residue-residue contacts also enable protein structure determination using metagenomic data [33] and structure prediction of homo- and heteromeric complexes [14, 43].

We tested whether *DomainRank* can improve residueresidue contact maps from different contact prediction methods. In total, we pooled 882 proteins from literature datasets and tested five different contact prediction algorithms. CCMpred [39], plm-DCA [6], and GREMLIN [20] are co-evolutionary contact prediction methods that analyze joint evolutionary couplings in multiple sequence alignments to predict contact maps. EPC-map combines evolutionary and physicochemical information for contact prediction [38]. MetaPSICOV uses a neural network to combine several co-evolutionary methods and further sequence-based features for contact prediction [18].

In this application domain, we used contact co-occurrence probabilities to construct the corroborating information graph. This co-occurrence probability reflects the likelihood that two contacts exist in an underlying secondary structure interface (Fig 3). We weight the edges in the corroborating information graph by the co-occurrence probability, which depends on the relative sequence shift of the contact and the secondary structure of the residues of the centered contact (the contact that is used as a reference point to assign co-occurrence probabilities to neighboring contacts).

**Figure 3:**
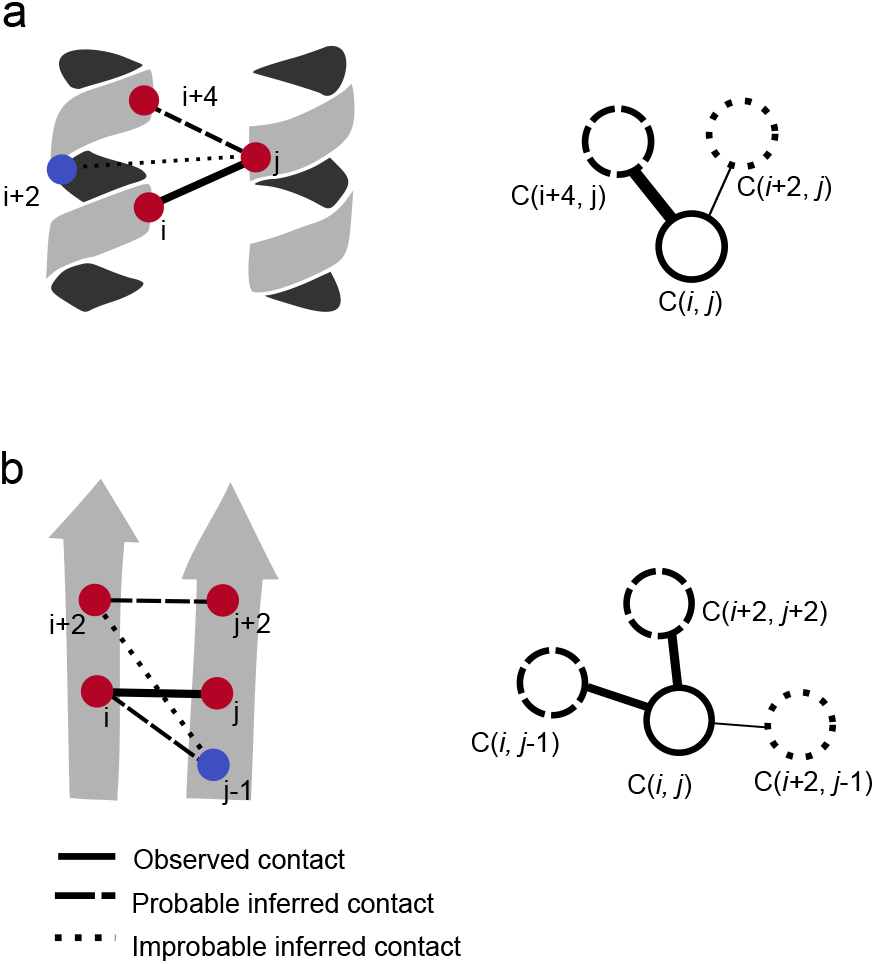
Contacts are embedded into the 3D protein structure and therefore occur in a pattern that is indicative for this structure. We illustrate this for a helix-helix and a strand-strand contact. (a) Consider a contact between i and j in two adjacent helices. The contact between i+2 and j is unlikely because the i+2 position is on the opposite side of the helix due to the helix geometry. In contrast, the contact between *i*+*4* and *j* is likely beause *i*+*4* is approximately one helix turn away from residue *i*+*4*. Thus, we connect the contacts *C*(*i, j*) and *C*(*i*+4, *j*) with ane dge in the corroborating information graph. (b) When observing the sheet-sheet contact *C*(*i, j*),there are likelycontactsbecauseofthe β-sheet geometry. In this example, *C*(*i*+2, *j*+2) and *C*(*i, j*−1) would be likely given the β-sheet structure while *C*(*i*+2, *j*−1) would be unlikely. Consequently, *C*(*i*+2, *j*−1) does not corroborate the observed contact *C*(*i, j*). Therefore, *C*(*i*+2, *j*−1) is not connected by an edge in the corroborating information graph.

For CCMpred, *DomainRank* improves the mean precision of the top *L/*2 and *L* contacts with sequence separation ≥24 by 10.4 and 10.8%, respectively (Fig 4a). The mean precision improvement of the other evolutionary contact prediction methods is similar: for plm-DCA the top *L/*2 and *L* precision improves by 11.6 and 6.7% (Fig 4b); for GREMLIN by 6.7 and 8.7% (Fig 4c). However, the precision improvement much smaller for EPC-map (2.9 and 3.0%, Fig 4d) and MetaPSICOV (0.9 and 1.2%, Fig 4e). The refinement results for contacts with sequence separation ≥ 12 residues are similar (S1 Fig).

**Figure 4:**
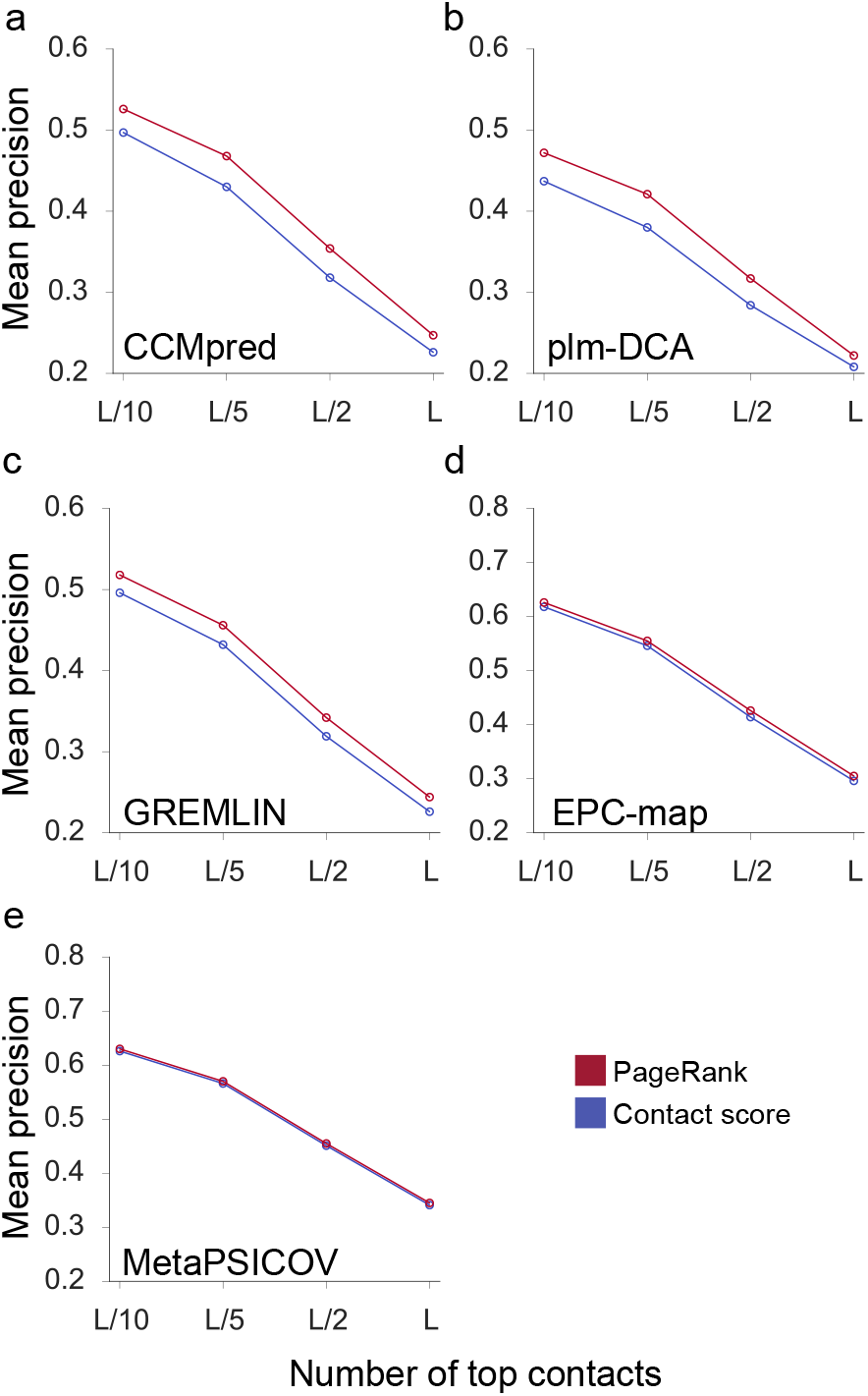
Contact refinement results using *DomainRank*. We refined the contact maps from the methods CCMpred, plm-DCA, GREMLIN, EPC-map, and MetaPSICOV. This figure shows the mean precision over 882 proteins pooled from different literature datasets. We show the results for the top ranked contacts with a sequence separation ≥24 residues. **a**: Comparison of CCMpred and *DomainRank* contacts. **b**: Comparison of plm-DCA and *DomainRank* contacts. **c**: Comparison of GREMLIN and *DomainRank* contacts. **d**: Comparison of EPC-map and *DomainRank* contacts. **e**: Comparison of MetaPSICOV and *DomainRank* contacts.

We attribute this to the kind of information that is already used by the input contact prediction methods. plm-DCA only uses evolutionary information, therefore the structural information about contact co-occurrence has the largest impact. EPC-map already uses structure information in the form of low-energy decoys. Thus, the contact maps by EPC-map are already more consistent with expected contact co-occurrence probabilities, and therefore the improvement of the contact maps is less pronounced. MetaPSICOV uses a two-stage classifier: The second classifier filters the first contact map by using local patches of predicted contacts [18]. Thus MetaPSICOV already uses contact co-occurrence probabilities to refine contact maps, albeit by training a neural network classifier. This might limit the applicability of our algorithm to contact prediction method that not already exploit an information source that are also leveraged by *DomainRank*.

Our approach is a simple post-processing method to increase contact map precision and can be easily added to any contact prediction pipeline with minimal effort.. Here too, we would like to emphasize that the contribution is in the improvement of data through *DomainRank*; we do not advance the state of the art in refining contact data.

### Refinement of Photo-Crosslinking/Mass Spectrometry Data

Crosslinking/mass spectrometry is a technique for studying the structure of proteins and protein complexes [35]. Crosslinkers are reagents that covalently link reactive residues in a protein if they are in spatial proximity. After digestion of the protein, the resulting peptides are subjected to mass spectrometric analysis. Specialized software matches crosslinked peptide pairs to the recorded mass spectra, which results in a list of scored crosslinks.

Our laboratory used the highly reactive photocrosslinker sulfo-SDA to increase crosslink density and used this data to reconstruct the domain structures of human serum albumin (HSA) with high resolution [2]. To the best of our knowledge, no attempts have yet been made to improve this kind of data with network-based algorithms. We tested whether *DomainRank* can increase the number of crosslinks at a given false discovery rate from a photo-crosslinking/mass spectrometry experiment (photo-CLMS). We performed crosslink searches with the search software xiSEARCH [11] and estimated the false discovery rate with a target-decoy method, which uses known wrong “decoy” sequences to estimate the error at a score cutoff [45, 27, 7].

The corroborating information in this application domain accounts for the reactivity of the used crosslinker sulfo-SDA. Sulfo-SDA contains an NHS-ester on one side and a diazirine on the other side. The activated diazirine group can react with any amino acid, which results in uncertain residue assignments (Fig 5). We assume that true-positive crosslinks are frequently supported by neighboring crosslink detections while decoy spectrum matches should rather appear in isolation (Fig 5a). Thus, crosslinks that appear in the same sequence region corroborate each other (Fig 5b) and common neighbors further reinforce this signal (Fig. 5c).

**Figure 5:**
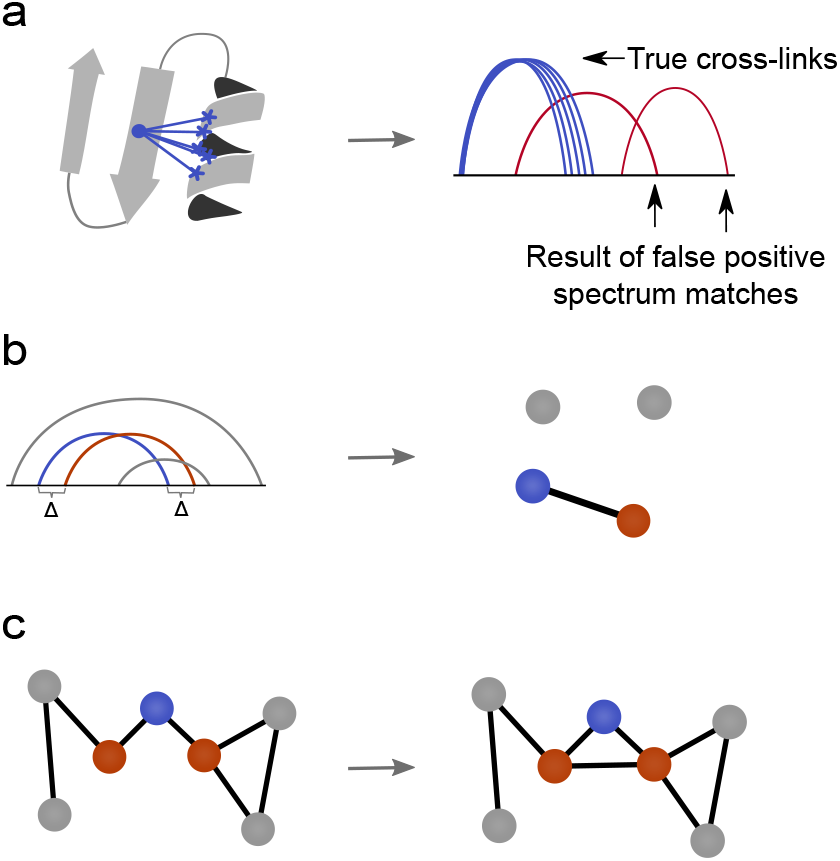
Corroborating evidence in CLMS data. (**a**) Because of the unspecific reactivity of sulfo-SDA, we expect to observe multiple neighboring crosslinks in a spatially close region of the protein. Thus, a high crosslink density in a region is indicative of true crosslinks. In contrast, crosslinks that are the result of wrong peptide spectrum matches should be isolated under the assumption that they are uniformly distributed over the protein sequence. (**b**) Two crosslinks are more likely to be correct if they link to the same residues within sequence separation Δ. In this figure, this condition holds for the blue and orange crosslink. Consequently, these crosslink nodes will be connected by an edge in the corroborating information graph. (**c**) Two crosslinks are more likely to be correct if they share a node in direct neighborhood in the corroborating information graph (transitivity). The crosslink nodes under consideration (orange) share one node in their neighborhood (blue). Due to transitivity, the red nodes will also be connected by an edge.

Fig 6a compares the HD-CLMS search results of xiSEARCH with the *DomainRank* post-processing on HSA. At 5% FDR, xiSEARCH identifies 397 crosslinks. At the same 5% FDR, *DomainRank* increases the number of crosslinks by 89.2% to 755 links. However, *DomainRank* does not increase the number at lower FDR values (such as 1% FDR). This is in line with our finding for protein-protein interactions: *DomainRank* effectively boosts low-confidence data while the effect on high-confidence data points is not as pronounced.

**Figure 6:**
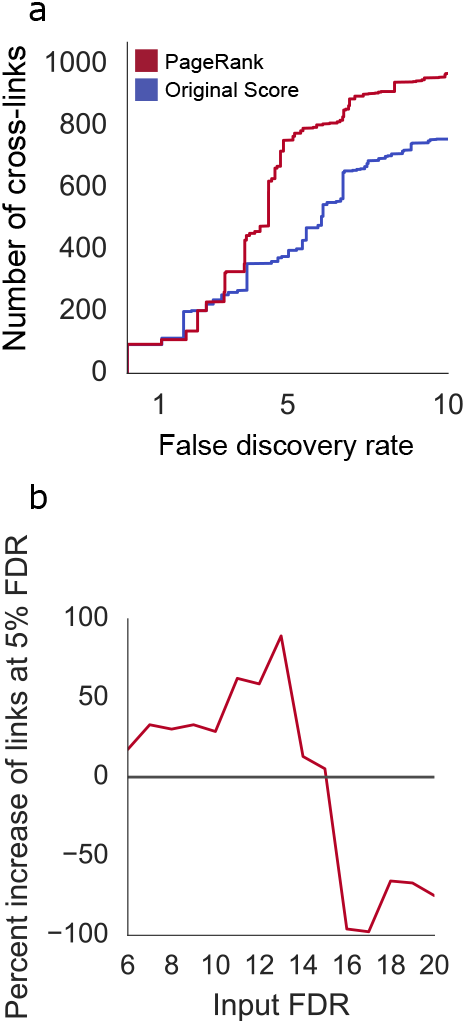
Corroborating evidence in photo-crosslinking/mass spectrometry data. (**a**) Number of crosslinks as a function of false discovery rate (FDR). At 5% FDR, xiSEARCH identifies 379 crosslinks while *DomainRank* identifies 755 crosslinks (89.2% increase). *DomainRank* does not increase the number of links at lower FDR values, indicating that especially low-confidence links are boosted. (**b**) Increase in number of identified links by *DomainRank* as function of input FDR. At 13% input FDR, most links are identified while the performance drops dramatically at higher FDR values. This suggests that too much noise in the input data deteriorates the performance of *DomainRank*.

We also tested the effect of the input FDR on the number of identified crosslinks (Fig 6b). From 6% to 13% input FDR, the number of links increases from 17.5% to 89.2%. However, higher input FDR data dramatically reduces the number of crosslinks (5.2% at 14% input FDR). With input FDR values of 15% and higher, the number of crosslinks decreases. This suggests that if too noisy data is used as input, the performance of *DomainRank* deteriorates. We recommend this type of analysis to find the optimal input FDR value for refining HD-CLMS data with *DomainRank*.

We also queried the crosslinks against the crystal structure of HSA (PDB ID 1AO6). For the 397 crosslinks identified by xiSEARCH at 5% FDR, 95.1% of the crosslinks are within 25 A in the crystal structure of HSA. 25 A is roughly the distance that a crosslink can bridge. For the 755 links identified by DomainRank at 5% FDR, 93.4% of the crosslinks are within 25 A in the crystal structure. This suggests that DomainRank broadly leads to the identification of crosslinks that have a similar agreement with crystal structure.

We showed that *DomainRank* dramatically increases the number of identified photo-crosslinks relative to the input search results from xiSEARCH at 5% FDR. And again, a final time, we state that our contribution lies in the relative improvement of data based on simple algorithmic template, rather than in an advancement of the field of crosslinking/mass spectromety.

## Conclusion

We presented *DomainRank*, a computational template based on PageRank, for improving biological interaction network data. The method consists of two steps: 1) Using corroborating information to embed the interaction data into a graph data structure. 2) Using Google’s PageRank algorithm to re-rank the interaction data. Interactions that are supported by corroborating information are ranked higher. We applied *DomainRank* to three biological network problems: on a protein-protein interaction network from quantitative proteomics experiments, on residue-residue contact pairs from evolutionary contact prediction methods, and on photo-crosslinking/mass spectrometry data. *DomainRank* improves the data in all three cases following the same, general template. Our results suggest that *DomainRank* is especially effective in boosting low-confidence interaction data.

While *DomainRank* is quite general, finding effective corroborating information might be the most time-consuming process. We view this analogous to the feature engineering process in machine learning that often determines the success of machine learning projects. This limitation aside, we think that our computational framework should be applicable to many other network datasets. We expect that for biological network projects, further performance gains will derive from more potent corroborating information. We think that this could be achieved by leveraging additional features by machine learning to predict edges or edge weights in the corroborating interaction graph. In summary, our proposed computational template can easily be applied in many application domains and allows researchers to implement their own algorithms to boost the confidence of their data with little effort.

## Acknowledgments

We thank Anders R. Kristensen for support on the quantitative mass spectrometry data analysis. We thank Kolja Stahl for proving residue-residue contact prediction results. This work was supported by the Wellcome Trust [103139, 10850] and DFG Grant [BR BR 2248/6-1, RA 2365/4-1]. The Wellcome Trust Centre for Cell Biology is supported by core funding from the Wellcome Trust [203149].

## Supplementary Matrial

### Applying *DomainRank* to Protein-Protein Interaction Data

#### Domain knowledge

Kristensen *et al*. [22] measured temporal changes in the interactome by a combination of quantitative proteomics and size exclusion chromatography. We used the original data set of the measured protein-protein interactions as a starting point. Our assumption is that two protein-protein interactions that occur in the same subcellular environment corroborate each other because the underlying biological mechanisms facilitate these interactions in this environment. We model this by computing a corroborating information score between protein-protein interaction pairs by computing the average agreement of subcellular localization GO-terms between the involved proteins.

#### Data set

We used measured protein-protein interactions from SEC-PCP-SILAC experiments by Kristensen et al. [22]. In this experiment, HeLa cells are labeled using amino acid isotopologs. Lysed cells are subjected to size exclusion chromatography and light isotope labeled proteins are used as standards. Proteins are identified by liquid-chromatography mass spectrometry and the light/medium ratios of the chromatograms are used to detect interactions, following the assumption that interacting proteins have similar elution profiles. We downloaded the protein identifications and chromatograms from the original study and used it as the input data for our experiments with *DomainRank* on protein-protein interaction data.

#### Data refinement

The input to refinement of protein-protein interaction data is a list of interactions from SEC-PCP-SILAC experiments. The initial scoring of the interactions is done by using the Euclidean distance (1-Euclidean distance/maximum Euclidean distance) with a maximum Euclidean distance cutoff of 6. We draw an edge in the corroborating information graph by overlap of subcellular localization GO-terms of the involved proteins. For any pair of proteins across interactions, we compute the overlap by the number of same GO-term annotations, normalized by the longer list of GO-terms of the two proteins. The GO-term score is the mean score of the protein pairs across interactions. We draw an edge if the score is higher or equal to 0.5.

The personalization vector **f** contains the Euclidean distance scores and we use the standard *DomainRank* damping parameter of α = 0.85.

#### Evaluation criteria

Kristensen et al. [22] found that Euclidean distance between chromatograms is the best predictor of interactions between two proteins in their experiment. Thus, we use the Euclidean distance (precisely, 1-Euclidean distance/maximum Euclidean distance) between chromatograms as a baseline, excluding low-quality chromatograms as described in the original manuscript. The list of interactions with Euclidean distances was used as an input to construct the corroborating information graph.

To evaluate the precision of the protein-protein interactions, we followed the following procedure: First, we took all proteins that were identified in the original study and also in the current version of the CORUM database of curated, mammalian protein-protein interactions [37]. Of all interactions containing these proteins, we considered all interactions that are also in the CORUM database as a true positive and as a false positive otherwise. For each of the individual three experiments of the original study, we select a cutoff for Euclidean distance and *DomainRank* score with a target precision score that is computed using CORUM interactions. We then pool the interactions of the three experiments and compute the final precision after removing redundant interactions that result from pooling of the three experiments. To perform a fair analysis on the same protein set, we use the intersection of proteins that are retained after sorting and selecting by Euclidean distance and *DomainRank* (and are also contained in CORUM). We then use all interactions above the chosen cutoff value to report the number of interactions.

#### Applying *DomainRank* to Contact Data

##### Domain knowledge

A common definition of residue– residue contacts is that their C_β_ atom distance is below 8 Å. Because the upper distance bound of a contact is tight, the distribution of contacts in the contact map should reflect the underlying regular structure of α-helices and β-sheets. Thus, a predicted contact map should agree with these regularities. Fig. 3 illustrates this idea for a contact between α-helices and β-strands. Given a helix-helix interface, contact *C*(*i, j*) and *C*(*i* + 2, *j*) contradict each other because the residue *i* + 2 points away from the helix interface (Fig. 3a). In contrast, contacts *C*(*i, j*) and *C*(*i* + 4, *j*) satisfy the physical constraints in α-helical structure. Thus, contacts *C*(*i, j*) and *C*(*i* + 4, *j*) corroborate each other and should have a large edge weight. We apply other patterns for different secondary structure pairs such as β-strands (Fig. 3b).

Instead of drawing binary edges between residue-residue interactions (contacts), we model the probability for the co-occurrence of two contacts by the probability of the relative sequence shift, conditioned on the secondary structure of the contacting residue pairs (denoted as *P*(*x, y*, Sec_1_, Sec_2_)). We obtain these probabilities from a training set of protein structures.

##### Data set

We used several datasets from literature and several contact prediction methods to validate *DomainRank*. We used validation datasets from SVMCON [4], PSICOV [17], ProC S3 [26], and MetaPSICOV [18]. Because we computed co-occurrence probability distributions on EPC-map training proteins, we filtered all proteins in the validation sets with an HHSearch [36] E-value below 10^*−*3^ to all proteins in the EPC-map training set [38]. This results in 882 with sizes from 25–499 residues. We used contact predictions from the following algorithms: plm-DCA [6], GREMLIN [20], CCMpred [39], EPC-map [38], and MetaPSICOV [18]. Plm-DCA, GREMLIN and CCMpred are evolutionary methods while EPC-map and MetaPSICOV are combination methods for contact prediction.

##### Data refinement

We construct the corroborating information graph by starting with the highest scoring interaction node (of contact *C*(*i, j*)) and edges to other contact nodes that are within eight positions around *C*(*i, j*) in the contact map. We weight the edges according to the probability distribution *P*(*x, y*, Sec_*i*_, Sec _*j*_), where Sec_*i*_ and Sec _*j*_ are the secondary structure assignments of residue *i* and *j*,as predicted by PSIPRED [16]. *x* and *y* denote the shift in sequence positions around *C*(*i, j*) of the second contact node.

We compute the probability distribution by analyzing the contacts of 742 proteins in the EPC-map training set [38]. We measure the co-occurrence frequencies of the relative shifts of the contacts, conditioned on the secondary structure of the centered contact. For distances between residue pairs slightly larger than 8 Å, we still consider this contact with a reduced count of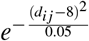. This reduces noise by considering residue pairs that are just out of the defined distance range and would not count otherwise. S2 Fig shows the co-occurrence probability matrices.

The personalization vector **f** is proportional to the scores from the input contact lists. In addition, we introduce a parameter β, which denotes the top β*L* contacts that we use to construct the corroborating information graph. Contact prediction methods might assign a probability to every possible contact. Therefore, the β parameter controls the number of contacts that are used in the algorithm, which would otherwise be the entire contact map. We tune the α and β parameters on 132 proteins from the EPC-map test dataset [38] which results in the following parameters: plm-DCA: α = 0.65, β = 2; GREMLIN: α = 0.6, β = 2.5; CCMpred: α = 0.45, β = 2; EPC-map: α = 0.35, β = 2.5; MetaPSICOV: α = 0.25, β = 2.5.

##### Evaluation criteria

A residue pair is defined to be in contact if their C_β_ atoms (C_α_ for glycine) are within 8 Å in the folded protein structure. We validate contact data for sequence separation cutoffs ≥12 and 24 residues separately. The higher the sequence separation cutoff, the more valuable are the contacts because they indicate non-local spatial interactions in the folded protein structure.For contact data, we also validate the top *L/*10, *L/*5, *L/*2, *L* contacts, where *L* is the number of amino acids of the protein.

As validation criteria, we use the precision TP/(TP+FP), where TP are true positives and FP are false positives.

## Applying *DomainRank* to CLMS Data

### Domain knowledge

A cross-link indicates that a residue pair is within a certain distance in the folded protein. The distance threshold is proportional to the crosslinker length. However, the mass spectrometric analysis of crosslinks, especially of photo crosslinking reagents, has technical limitations that introduces noise into the measurement. These noisy crosslinks are not within crosslink distance in the native structure.

We now introduce the corroborating information in crosslink data that we use to reduce noise and therefore increase the number of crosslinks of residue pairs that are within 25 Å in the native structure.

Since one side of the sulfo-SDA crosslinker is unspecific, this side should react with multiple neighboring residues that are spatially close in independent crosslink events. In these high-density crosslink regions, true crosslinks are frequently supported by neighboring crosslink detections. In contrast, crosslinks that result from decoy peptide spectrum matches (target-decoy or decoy-decoy) should rather appear in isolation. This assumes that the probability of observing a decoy spectrum match is uniform along the protein chain (Fig 5a). Thus, target crosslink matches in dense regions corroborate each other. We include this information by connecting two crosslink nodes if the crosslinks connect he same sequence region within a sequence separation Δ (Fig 5b). This also accounts for uncertain assignments of the crosslinked residues. We further reinforce this signal by connecting crosslink nodes that share a common neighbor (Fig 5c). This forms dense connections in regions with high corroborating information which increases the propagation of information in that region.

### Dataset

To demonstrate our method on HD-CLMS data, we used the dataset from Belsom *et al*. [2]. This study crosslinked human serum albumin using sulfo-SDA. This results in 399/795 identified links at 5/10% FDR. Note that the crosslink searches in this work have been recalculated using a newer version of the search software xiSEARCH [10] and therefore deviate slightly from the original study.

### Data refinement

The input to HD-CLMS data refinement is a list of crosslink and decoy matches, sorted by the score of the search algorithm xiSEARCH [11].

A node in the corroborating information graph represents a crosslink. We then connect a node by an edge if the crosslinks are within Δ = 6 sequence separation in both peptides. Then, we connect all nodes that share at least one neighbor in the graph. We repeat this process two times.

Note that xiSEARCH uses inverted input sequences for FDR calculation to represent decoys. Therefore, we map residue positions in decoy sequences back to the original target sequence space. This ensures that target hits and decoy hits are mapped to the same space when we construct the corroborating information graph.

The personalization vector **f** contains probabilities that are proportional to the crosslink scores. We use the standard damping parameter of α = 0.85.

### Evaluation criteria

We evaluate the re-scoring of HD-CLMS data by evaluating the yield of crosslinks at a predefined false discovery rate (FDR) [45, 7]. We calculate the false discovery rate by first summarizing all peptide-spectrum matches to peptide pairs by taking the squared sum of their scores from xiSEARCH [10]. Peptide pairs are summarized to links by the same procedure. For all links that pass a certain score cutoff, we compute the FDR by the following formula:

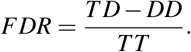

Here, *TD* is a target-decoy match (having one target and one decoy sequence), *DD* is a decoy-decoy match, and *TT* is a target-target match. We report the number of target-target hits at a pre-defined FDR (5% in this work).

